# Activation of GSDME compensates for GSDMD deficiency in a mouse model of NLRP3 inflammasomopathy

**DOI:** 10.1101/2021.01.06.425634

**Authors:** Chun Wang, Tong Yang, Jianqiu Xiao, Canxin Xu, Yael Alippe, Kai Sun, Thirumala-Devi Kanneganti, Joseph B. Monahan, Yousef Abu-Amer, Judy Lieberman, Gabriel Mbalaviele

**Author notes:** To whom correspondence should be addressed. Division of Bone and Mineral Diseases, Washington University School of Medicine, 660 South Euclid Avenue, Campus Box 8301, St. Louis, MO 63110. These authors contributed equally to this work (Chun Wang, Tong Yang, Jianqiu Xiao).

## Abstract

The D301N NLRP3 mutation in mice (D303N in humans) causes severe multi-organ damage and early death driven by the constitutively activated NLRP3 (NLRP3^ca^) inflammasome. Triggered inflammasomes activate caspase-1 to process IL-1 family cytokines and gasdermin D (GSDMD), generating N-terminal fragments, which oligomerize within the plasma membrane to form pores, which cause inflammatory cell death (pyroptosis) and through which IL-1β and IL-18 are secreted. GSDMD activation is central to disease symptoms since spontaneous inflammation in *Nlrp3*^*ca*^;*Gsdmd*^*-/-*^ mice is negligible. Unexpectedly, when *Nlrp3*^*ca*^;*Gsdmd*^*-/-*^ mice were challenged with LPS or TNF-α, they secreted high amounts of IL-1β and IL-18, suggesting an alternative GSDMD-independent inflammatory pathway. Here we show that GSDMD deficient macrophages subjected to inflammatory stimuli activate caspase-8, -3 and GSDME-dependent cytokine release and pyroptosis. Caspase-8, -3 and GSDME also activated pyroptosis when NLRP3 was stimulated in caspase-1 deficient macrophages. Thus, a salvage caspase-8, -3-GSDME inflammatory pathway is activated following NLRP3 activation when the canonical NLRP3-caspase-1-GSDMD is blocked. Surprisingly, the active metabolite of the GSDMD-inhibitor disulfiram, inhibited not only GSDMD but also GSDME-mediated inflammation *in vitro* and suppressed severe inflammatory disease symptoms in *Nlrp3*^*ca*^ mice, a model for severe neonatal multisystem inflammatory disease. Although disulfiram did not directly inhibit GSDME, it suppressed inflammasome activation in GSDMD-deficient cells. Thus, the combination of inflammatory signals and NLRP3^ca^ overwhelmed the protection provided by GSDMD deficiency, rewiring signaling cascades through caspase-8, -3 and GSDME to propagate inflammation. This functional redundancy suggests that concomitant inhibition of GSDMD and GSDME may be necessary to suppress disease in inflammasomopathy patients.

## Introduction

Upon sensing plasma membrane perturbations caused by certain microbial products or sterile danger signals, the innate immune receptor NOD-like receptor (NLR) family, pyrin domain containing 3 (NLRP3) assembles an intracellular protein complex known as the NLRP3 inflammasome ^1, 2, 3^. This macromolecular structure is formed by NLRP3, the adaptor protein apoptosis-associated speck-like protein containing a CARD (ASC), and caspase-1, which becomes activated when it is recruited to inflammasomes. Caspase-1, which is activated by all the canonical inflammasomes, catalyzes the maturation of interleukin-1β (IL-1β) and IL-18 ^2, 3^, and also processes gasdermin D (GSDMD), generating N-terminal fragments, which oligomerize within the plasma membrane to form pores through which IL-1β and IL-18 are secreted ^4, 5, 6, 7, 8^. Excessive pore formation compromises the integrity of the plasma membrane, causing a lytic form of cell death known as pyroptosis. After some signals, cells survive GSDMD activation, but still process and release inflammatory cytokines in a GSDMD-dependent process and continue their other functions, and hence have been termed “hyperactivated” ^6^. For the whole organism, pyroptosis is a double-edged sword - it releases inflammatory factors that recruit immune cells to sites of pathogenic infection, which attack and phagocytose pathogens, destroy their replication niches and boost adaptive immune responses ^9, 10^. However, excessive or uncontrolled pyroptosis and inflammatory cytokine release can lead to organ damage, circulatory collapse or even death ^11, 12, 13^.

GSDMD is also cleaved by caspase-11 (mouse ortholog of human caspase-4 and -5) ^14^, neutrophil elastase and cathepsin G ^15, 16, 17 18^, caspase-3 ^19^, and caspase-8 ^20, 21^. Cleavage by all of these proteases, except caspase-3, which inactivates GSDMD, produces a pore-forming N-terminal fragment. Caspase-3 also processes gasdermin E (GSDME, also known as DNFA5), producing active N-terminal fragments whose pore-forming activity promotes pyroptosis in response to apoptotic stimuli ^22, 23, 24^, including chemotherapeutic drugs ^25, 26^, *Yersinia* infection ^20^, glucocorticoid treatment ^24^, and inflammasome activators in cells lacking GSDMD or caspase-1 ^27, 28^. GSDME is also a substrate of granzyme B ^23, 29^. The role of GSDME in cell death may be cell context-dependent as it is seemingly dispensable for secondary necrosis in certain cell types ^30, 31^. Its function in mediating the release of inflammatory cytokines also remains unclear. A recent study reported that IL-1α was secreted through GSDME conduits in caspase-1 and -11 deficient macrophages, but pro-IL-1β was neither processed nor released ^28^.

Chronic activation of the NLRP3 inflammasome by endogenous host danger-associated molecular patterns and the ensuing excessive production of IL-1β and IL-18 underlie the pathogenesis of several autoimmune and autoinflammatory diseases, including inflammatory bowel disease ^32, 33, 34^, and rheumatoid arthritis ^35, 36, 37^, and contribute to the severity of atherosclerosis ^38, 39^, gout ^40, 41, 42, 43^, and diabetes ^44^. Gain-of-function mutations in *NLRP3* also cause a spectrum of autoinflammatory disorders known as cryopyrin-associated periodic syndromes (CAPS), whose severity is linked to specific *NLRP3* mutations. Neonatal-onset multisystem inflammatory disease (NOMID) is the most severe and familial cold autoinflammatory syndrome (FCAS) and Muckle-Wells syndrome (MWS) are less severe ^12, 13^. Clinical manifestations of CAPS include systemic inflammation, skin lesions, central nervous system symptoms including headache, hearing loss and learning difficulties, and skeletal anomalies ^13, 45, 46, 47, 48, 49^. Mice genetically engineered to express NLRP3 variants that harbor mutations found in CAPS patients (NLRP3^ca^) develop severe systemic inflammation characterized by excessive secretion of IL-1β and IL-18, multi-organ damage, and premature death ^50, 51^. Although young NOMID mice lacking the IL-1 receptor do not develop inflammation ^52^, a persistent low-grade inflammation is reported in FCAS mice and MWS mice with defective IL-1 and IL-18 signaling ^53^, suggesting that pyroptosis may be the culprit. In support of this view, knocking out GSDMD prevents the pathogenesis of NOMID ^54^, Familial Mediterranean Fever (FMF) ^55^, and experimental autoimmune encephalitis ^56^. However, how these mutant mice with an underlying dysregulated NLRP3 inflammasome activity withstand superimposed inflammatory challenges has not been studied.

CAPS and several other diseases of dysregulated inflammasome activity, including FMF and macrophage activation syndrome, which are caused by *PYRIN* and *NLRC4* mutations, respectively, are treated with IL-1 blockers ^57^. These therapies have significantly improved the quality of life of affected individuals, but the disease does not resolve in some patients ^58, 59, 60, 61^. The underlying mechanisms of resistance are not understood, but lingering low-grade inflammation driven by IL-18 and pyroptosis is suspected, as noted above. To identify drugs that block the integrating nodes of inflammasome signaling, recent studies screened several safe marketed drugs for anti-GSDMD activity. They identified disulfiram, also known as Antabuse, an FDA-approved drug for the treatment of alcohol addiction, and dimethyl fumarate, also known as Tecfidera used to treat multiple sclerosis, as inhibitors of GSDMD and pyroptosis ^56, 62^. Both drugs also inhibited LPS-induced IL-1β and IL-18 secretion *in vitro* and *in vivo* ^63^. These observations provide a rationale for evaluating pyroptosis inhibitors in animal inflammasomopathy models.

In this study, we challenged mice bearing dysregulated constitutively active *Nlrp3* mutations (*Nlrp3*^*ca*^) with inflammatory stimuli (LPS or TNF-α) to understand better the role of GSDMD in promoting inflammation. We found that these intense NLRP3 inflammasome-activating signals overwhelmed the protection afforded by *Gsdmd* ablation and caused pyroptosis and pro-inflammatory IL-1 family cytokine processing and secretion in *Nlrp3*^*ca*^*;Gsdmd*^*-/-*^ macrophages. Unexpectedly, these stimuli activated caspase-8, -3 and GSDME, which were responsible for pyroptosis and pro-inflammatory cytokine release. We also found that the active metabolite of disulfiram inhibited the functions of both GSDMD and GSDME, and protected mice from the pathology caused by dysregulated NLRP3 inflammasome.

## Results

### GSDMD deficient mice expressing hyperactive NLRP3 inflammasome age normally

D301N substitution in NLRP3 imposes conformational changes and constitutive activation of NLRP3 (*Nlrp3*^*ca*^), ultimately causing an inflammatory disease in mice that resembles human NOMID ^50, 64^. Most *Nlrp3*^*ca*^ mice die prematurely around 3 weeks of age due to severe systemic inflammation that damages multiple organs, including the skin, brain, bones, and spleen ^50, 64, 65^. *Gsdmd*^*-/-*^ and *Nlrp3*^*ca*^ mice lacking GSDMD (*Nlrp3*^*ca*^;*Gsdmd*^*-/-*^ mice) grew normally and were indistinguishable from their wild-type (WT) counterparts when monitored for up to 66 days, indicating strong GSDMD-dependence of inflammation in NOMID mice ^54^. However, mice expressing constitutively activated NLRP3 as a result of L351P substitution that were genetically deficient in *Il-1* and *Il-18* unexpectedly showed signs of lingering inflammation ^53^. This finding prompted us to monitor *Nlrp3*^*ca*^;*Gsdmd*^*-/-*^ mice for longer. Mouse survival, white blood cell (WBC) counts, and spleen weight of *Nlrp3*^*ca*^;*Gsdmd*^*-/-*^ and *Gsdmd*^*-/-*^ mice remained indistinguishable at 6 months and 12 months of age (Fig. S1A). Accordingly, the architecture of the spleen and the liver was histologically similar between both genotypes (Fig. S1B and C). Thus, *Nlrp3*^*ca*^;*Gsdmd*^*-/-*^ mice age normally in homeostatic conditions, consistent with our previous study ^54^.

### LPS and TNF-α stimulate IL-1β and IL-18 secretion not only in *Nlrp3*^*ca*^ but also *Nlrp3*^*ca*^;*Gsdmd*^*-/-*^ mice

*Gsdmd*^*-/-*^ mice, unlike *caspase-11*^*-/-*^ mice, were not fully protected from death caused by a lethal dose of LPS ^14^, suggesting that there might be a GSDMD-independent inflammatory pathway. Baseline serum IL-1β, IL-18 and IL-6 were low and comparable in WT and *Gsdmd*^*-/-*^ mice but were constitutively increased in *Nlrp3*^*ca*^ mice and, as expected, returned to normal in *Nlrp3*^*ca*^;*Gsdmd*^*-/-*^ mice (Fig. 1A). After LPS challenge, serum IL-1β and IL-18 increased in WT mice, but not in *Gsdmd*^*-/-*^ mice. However, in *Nlrp3*^*ca*^ mice these cytokine levels were about 10-fold higher than in LPS-stimulated WT mice and surprisingly were not substantially reduced in *Nlrp3*^*ca*^;*Gsdmd*^*-/-*^ mice. LPS stimulated high serum levels of IL-6 and CXCL1 in all the genotypes. TNF-α injection increased serum IL-1β and IL-18 in *Nlrp3*^*ca*^, but not in WT or *Gsdmd*^*-/-*^ mice, but this increase was significantly reduced but not eliminated in *Nlrp3*^*ca*^;*Gsdmd*^*-/-*^ mice. TNF-α induced elevated IL-6 and CXCL1 in all genotypes, but induced more IL-6 in *Nlrp3*^*ca*^ mice, which was still above the level in WT mice. These results indicate that both GSDMD-dependent and GSDMD-independent inflammation is induced in the setting of constitutive NLRP3 activation.

**Fig 1.**
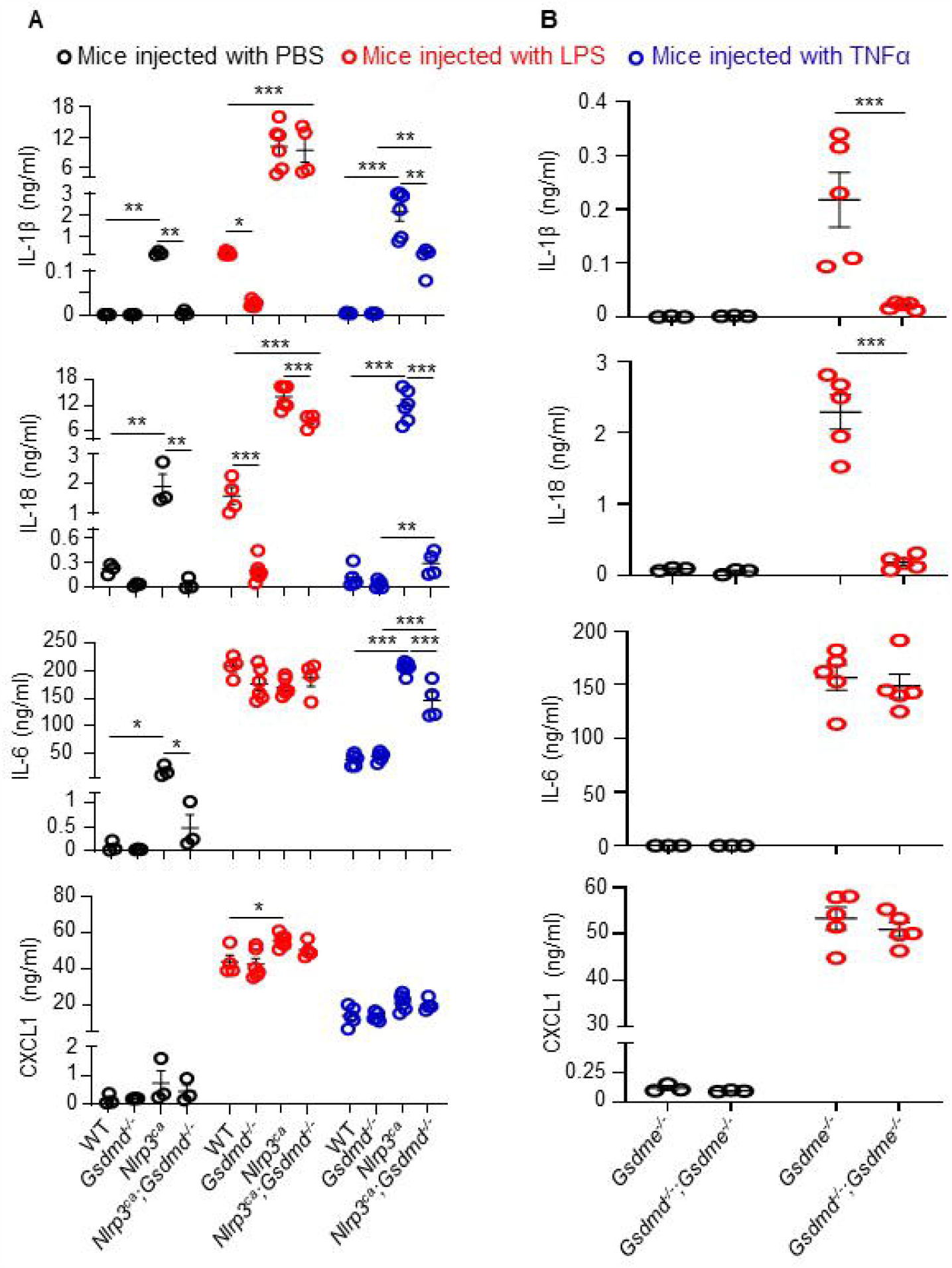
LPS and TNF-α induce the production of cytokines and chemokines even in *Nlrp3*^*ca*^;*Gsdmd*^*-/-*^ mice. (A) Three-month-old WT, *Gsdmd*^*−/−*^, *Nlrp3*^*ca*^, and *Nlrp3*^*ca*^;*Gsdmd*^*−/−*^ mice were injected with 15 mg/kg LPS for 6 hours or 0.5 mg/kg TNF-α for 2 hours. (B) Three-month-old *Gsdme*^*−/−*^ *and Gsdmd*^*−/−*^;*Gsdme*^*−/−*^ mice were injected with 15 mg/kg LPS for 6 hours. PBS-administrated mice served as controls. Serum cytokine levels were measured by V-PLEX Plus Proinflammatory Panel 1 Mouse Kit, except for IL-18, which were assessed by ELISA. Data are mean ± SEM. *P < 0.05; **P < 0.01; ***P < 0.001. LPS, lipopolysaccharide; IL, interleukin; WT, wild type; ca, constitutive activation; TNF-α, tumor necrosis factor α.

To determine whether GSDMD-independent inflammation in LPS-treated mice might be due to GSDME activation, we compared baseline and LPS-stimulated serum cytokines (IL-1β, IL-18, IL-6, and CXCL1) in *Gsdme*^*-/-*^ and *Gsdmd*^*-/-*^*;Gsdme*^*-/-*^ animals bearing unmutated *Nlrp3* (Fig. 1B). *Gsdme* deficiency did not cause baseline cytokine elevation. After LPS challenge, the IL-1 family cytokines were similarly elevated in *Gsdme*^*-/-*^ mice as in WT mice but did not increase in *Gsdmd*^*-/-*^*;Gsdme*^*-/-*^ animals. LPS and TNF-α induced elevated IL-6 and CXCL1 in all genotypes. These results suggest that GSDME does not play a role in IL-1β and IL-18 secretion in GSDMD sufficient mice that do not expressed constitutively active NLRP3.

### Macrophages lacking GSDMD secrete IL-1β through GSDME

To study NLRP3 inflammasome signaling in GSDMD sufficient or insufficient cells, we treated WT, *Nlrp*3^*ca*^, and *Nlrp*3^*ca*^*;Gsdmd*^*-/-*^ bone marrow-derived macrophages (BMDMs) with LPS in the absence or presence of nigericin, an activator of the NLRP3 inflammasome ^66^. While the combination of LPS and nigericin was required for the maturation of GSDMD and caspase-1 in WT BMDMs, endotoxin alone induced these responses in *Nlrp*3^*ca*^ and *Nlrp*3^*ca*^*;Gsdmd*^*-/-*^ cells (Fig. 2A), consistent with the constitutively activated state of this mutated protein ^50^. However, the combination of LPS and nigericin caused more GSDMD and caspase-1 cleavage than LPS alone in cells with constitutively active NLRP3. In these cells whenever caspase-1 was cleaved, there was a faint caspase-8, -3 cleavage band. Although full length GSDME was easily visualized in WT and *Nlrp*3^*ca*^ BMDMs, cleavage of GSDME was not detected. However, in *Nlrp*3^*ca*^*;Gsdmd*^*-/-*^ cells, although basal levels of caspase-8, -3 and GSDME appeared similar to levels in WT or *Nlrp*3^*ca*^ BMDMs, cleaved caspase-8 (p18 fragment) and caspase-3 (p17 fragment) were more prominent and GSDME cleavage was readily detected (Fig. 2A). Neither caspase-3 nor GSDME cleavage in *Nlrp*3^*ca*^*;Gsdmd*^*-/-*^ BMDMs required nigericin. Thus, in *Nlrp*3^*ca*^*;Gsdmd*^*-/-*^ cells, caspase-8, -3 and GSDME are robustly activated by LPS.

**Fig 2.**
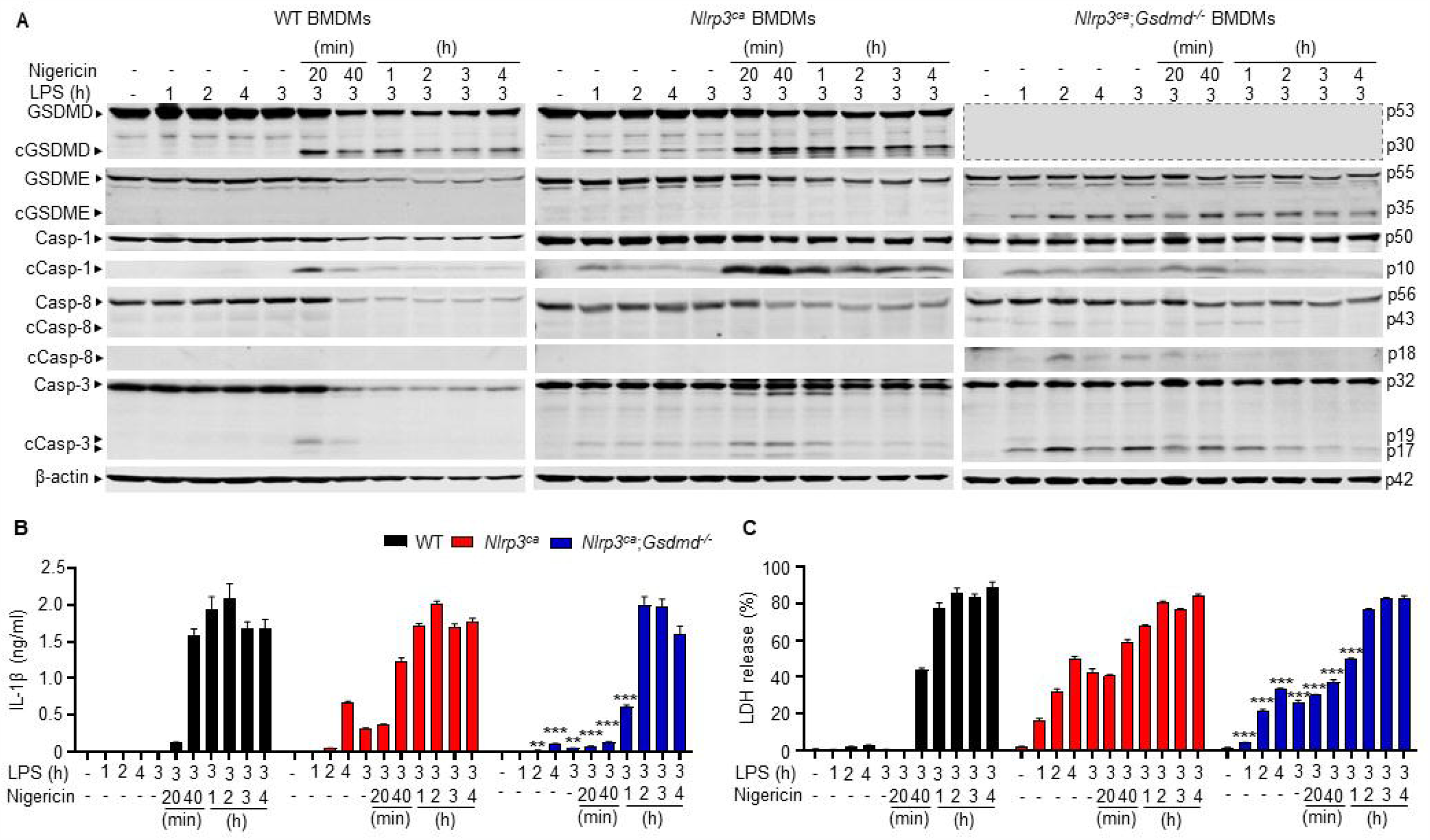
LPS and nigericin induce the release of IL-1β and LDH by *Nlrp3*^*ca*^;*Gsdmd*^*−/−*^ BMDMs. BMDMs were expanded *in vitro* in M-CSF-containing media from bone marrow cells isolated from WT, *Nlrp3*^*ca*^, or *Nlrp3*^*ca*^;*Gsdmd*^*−/−*^ mice. BMDMs were primed with 100 ng/ml LPS for 1, 2, 3, or 4 hours and treated with nigericin for 20 minutes, 40 minutes, 1, 2, 3, 4 hours. (A) Proteins in the whole cell lysates were analyzed by immunoblotting. The dash lined box was drawn to fill the void. IL-1β (B) and LDH (C) in the conditioned media were measured by ELISA and by the cytotoxicity detection Kit, respectively. Data are mean ± SEM from experimental triplicates and are representative of at least three independent experiments. **P < 0.01; ***P < 0.001 vs corresponding data in *Nlrp3*^*ca*^. BMDMs, bone marrow-derived macrophages; cCasp, cleaved caspase; cGSDM, cleaved gasdermin; h, hour; IL-1β, interleukin-1β; LDH, lactate dehydrogenase; LPS, lipopolysaccharide; min, minute; M-CSF, macrophage colony-stimulating factor; WT, wild type; ca, constitutive activation.

To examine how GSDMD deficiency affects IL-1β secretion, its levels were measured after nigericin treatment of LPS primed WT, *Nlrp3*^*ca*^, or *Nlrp3*^*ca*^;*Gsdmd*^*-/-*^ BMDMs. LPS alone induced IL-1β secretion in *Nlrp3*^*ca*^ cells (Fig. 2B). Nigericin increased the release of IL-1β by LPS-primed cells in the three genotypes, though this response was delayed in double mutant cells, reaching levels comparable to WT and *Nlrp3*^*ca*^ cells by 2 hours. We also assessed cell death by measuring the release of lactate dehydrogenase (LDH). While LPS and nigericin were required for LDH release in WT BMDMs, endotoxin alone induced these responses in *Nlrp*3^*ca*^ and *Nlrp*3^*ca*^*;Gsdmd*^*-/-*^ cells, though this response was attenuated in double mutant cells and enhanced by nigericin in both genotypes (Fig. 2C). Thus, activated *Nlrp*3^*ca*^*;Gsdmd*^*-/-*^ BMDMs secreted IL-1β, and underwent pyroptosis, outcomes that correlated with GSDME maturation.

To examine further the impact of GSDMD and GSDME deficiency on LPS plus nigericin induced pyroptosis and IL-1β processing and release, WT, *Gsdmd*^*-/-*^, *Gsdme*^*-/-*^, and *Gsdmd*^*-/-*^ *;Gsdme*^*-/-*^ BMDMs lacking constitutively activated NLRP3 were pretreated with LPS for 3 hours and then with nigericin for up to 4 hours. Within 20 minutes of adding nigericin, caspase-1, and -<u>3</u> were cleaved in all 4 cell lines (Fig. 3A; Fig. S2A and B). Similar to our findings in Fig. 2A in *Nlrp3*^*ca*^ BMDMs, in cells bearing unmutated *Nlrp3* treated with LPS and nigericin, GSDMD was cleaved in WT and *Gsdme*^*-/-*^ BMDMs, and caspase-8 p18 and caspase-3 p17 fragment were more prominent in cells lacking GSDMD. GSDME cleavage was detected only in *Gsdmd*^*-/-*^ cells. GSDMD and GSDME cleavage was detected at the same time as caspase activation. Since GSDME was not cleaved in WT BMDMs, the time-dependent decline of its levels in WT cells was likely secondary to pyroptosis. Thus, GSDME cleavage occurs after NLRP3 inflammasome activation in GSDMD-deficient BMDMs.

**Fig 3.**
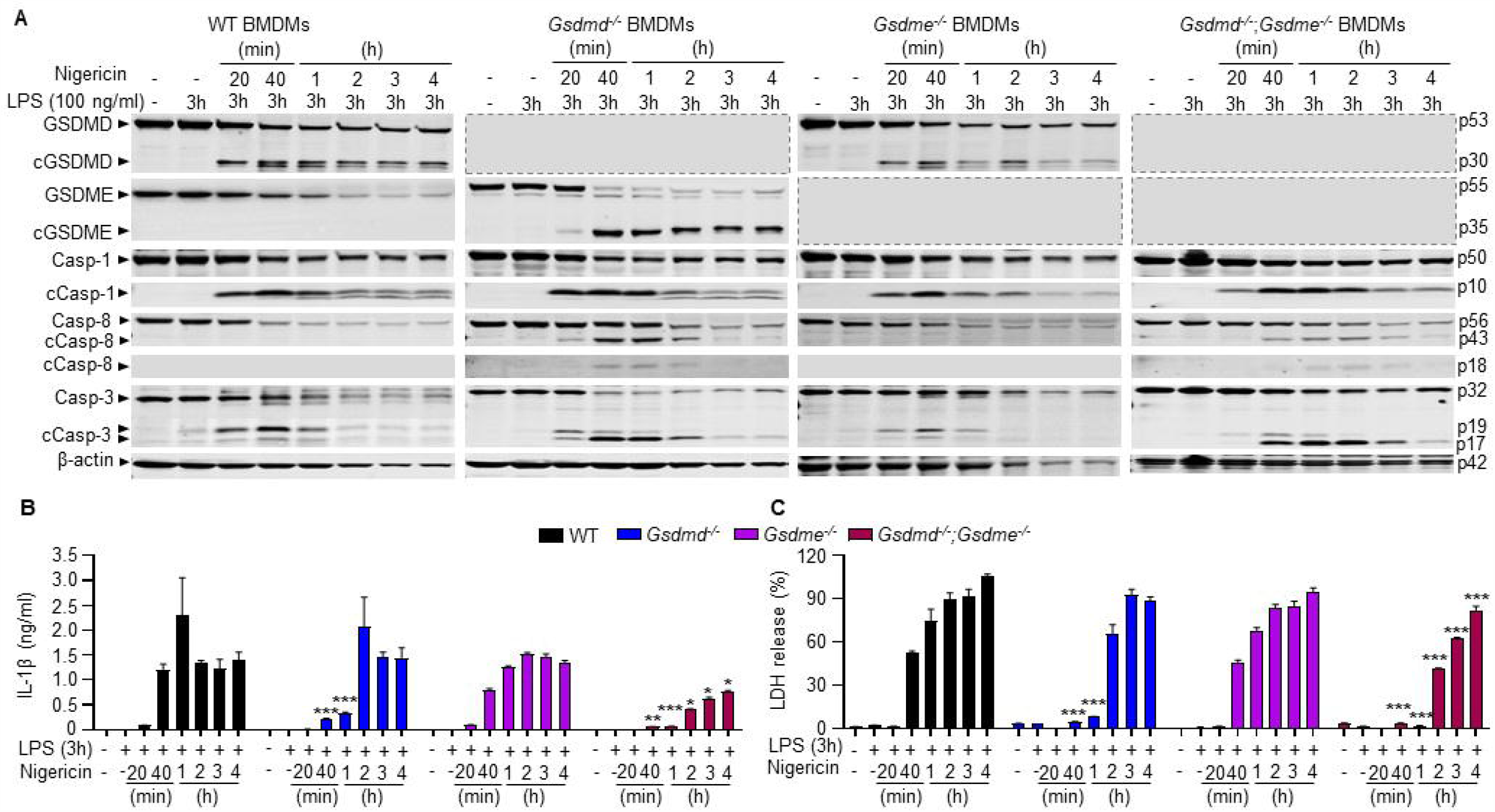
GSDME is involved in the release of IL-1β and LDH induced by LPS and nigericin. BMDMs were expanded *in vitro* M-CSF-containing media from bone marrow cells isolated from WT, *Gsdmd*^*−/−*^, *Gsdme*^*−/−*^, or *Gsdmd*^*−/−*^;*Gsdme*^*−/−*^ mice. BMDMs were primed with 100 ng/ml LPS for 3 hours and treated with nigericin for 20 minutes, 40 minutes, 1, 2, 3, 4 hours. (A) Proteins in the whole cell lysates were analyzed by immunoblotting. Dash lined boxes were drawn to fill the void. IL-1β (B) and LDH (C) in the conditioned media were measured by ELISA and by the cytotoxicity detection Kit, respectively. Data are mean ± SEM from experimental triplicates and are representative of at least three independent experiments. *P < 0.05; **P < 0.01; ***P < 0.001 vs corresponding data in WT. BMDMs, bone marrow-derived macrophages; cCasp, cleaved caspase; cGSDM, cleaved gasdermin; h, hour; IL-1β, interleukin-1β; LDH, lactate dehydrogenase; LPS, lipopolysaccharide; min, minute; nig, nigericin; M-CSF, macrophage colony-stimulating factor; WT, wild type.

Next, we examined whether caspase-1 was required to activate caspase-3 and GSDME by treating caspase-1 deficient BMDMs with LPS and nigericin. Consistent with previous reports ^67^, LPS and nigericin activated caspase-3, and promoted the generation of the GSDMD p10 fragment in *Casp1*^*-/-*^ BMDMs, although caspase-3 and GSDMD processing were delayed compared to caspase-1 sufficient cells (Fig. S2C). GSDME cleavage was detected in *Casp1*^*-/-*^ BMDMs at the same time as fully processed caspase-3. Thus, although the maturation of caspase-1 and GSDMD is unperturbed by GSDME deficiency, NLRP3 inflammasome activation forces GSDME processing in cells lacking either caspase-1 or GSDMD.

We examined the extent to which GSDME mediates IL-1β and LDH release in GSDMD deficient cells. Although LPS-primed *Gsdmd*^*-/-*^ BMDMs did not release IL-1β or LDH at early time-points up to 1 hour after nigericin addition, they were as active as WT cells in secreting IL-1β and releasing LDH at later time-points (Fig. 3B and C) and in dose-dependent manner (Fig. S2D). Loss of GSDME did not affect IL-1β and pyroptosis, but these readouts were significantly impaired, but not abrogated, in *Gsdmd*^*-/-*^;*Gsdme*^*-/-*^ BMDMs (Fig. 3B). Likewise, IL-1β processing was blunted, and LDH release was delayed in *caspase-1*^*-/-*^ BMDMs (Fig. S2E). Collectively, these data suggest that the NLRP3 inflammasome stimuli force IL-1β secretion and pyroptosis through GSDME in GSDMD or caspase-1 deficient cells.

### A disulfiram metabolite inhibits both GSDMD-dependent and -independent pyroptosis

Disulfiram and its active metabolite, bis(diethyldithiocarbamate)-copper (CuET) antagonize GSDMD pore-forming activity but have not been reported to inhibit GSDME pore formation ^62^. To investigate whether these compounds might inhibit GSDMD-independent pyroptosis, WT, *Gsdmd*^*-/-*^, *Gsdme*^*-/-*^, and *Gsdmd*^*-/-*^;*Gsdme*^*-/-*^ BMDMs were treated with LPS for 3 hours and then CuET or vehicle for 1 hour followed by an additional 3 hours of incubation with nigericin. In the absence of CuET, GSDMD was cleaved in WT and *Gsdme*^*-/-*^ BMDMs (Fig. 4A) and caspase-1 deficient BMDMs (Fig. S3A), consistent with the results shown in Fig. 3. While GSDME was cleaved only in *Gsdmd*^*-/-*^ cells, caspase-1 and caspase-3 were cleaved in all cell lines. However, although the full-length caspase protein bands sharply decreased, the signals of their cleaved fragments were faint (Fig. 4A), presumably because of pyroptosis, which was maximal in cells treated with nigericin for 2 hours. GSDMD, GSDME, and caspase-1 cleavage was consistently inhibited by CuET, in a dose-dependent manner (Fig. 4A; Fig. S3A). The band for cleaved caspase-3 was more prominent in CuET-treated than untreated WT, *Gsdmd*^*-/-*^, and *Gsdme*^*-/-*^ BMDMs, which may have been the result of attenuated cell death. CuET inhibited the release of IL-1β and LDH in all cell genotypes, but was less efficacious in *Gsdmd*^*-/-*^;*Gsdme*^*-/-*^ BMDMs, in which cytokine release and pyroptosis were attenuated (Fig. 4B and C; Fig. S3B). Disulfiram and CuET both inhibited IL-1β and LDH release in WT BMDMs with comparable dose response curves (Fig. S3C-F). Thus, CuET inhibits the maturation of both GSDMD and GSDME and consequently, inhibits both GSDMD-dependent and -independent pyroptosis and IL-1β release.

**Fig 4.**
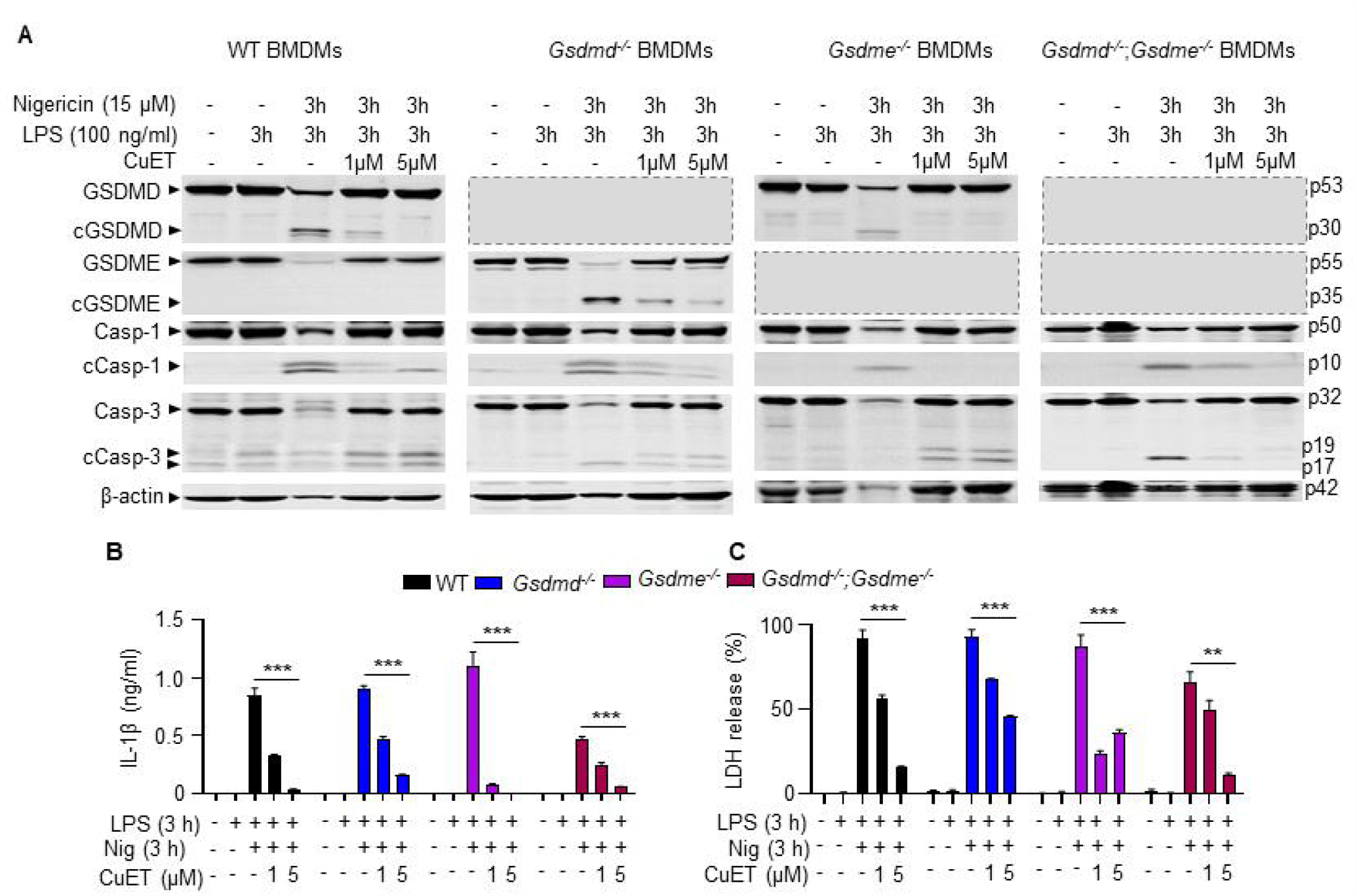
CuET inhibits NLRP3 inflammasome-dependent processing of GSDMs and caspases. BMDMs were expanded *in vitro* M-CSF-containing media from bone marrow cells isolated from WT, *Gsdmd*^*−/−*^, *Gsdme*^*−/−*^, or *Gsdmd*^*−/−*^;*Gsdme*^*−/−*^ mice. BMDMs were primed with 100 ng/ml LPS for 3 hours and treated with vehicle or CuET for 1 hour before adding 15 µM nigericin for 3 hours. (A) Proteins in the whole cell lysates were analyzed by immunoblotting. IL-1β (B) and LDH (C) in the conditioned media were measured by ELISA and by the cytotoxicity detection Kit, respectively. Data are mean ± SEM from experimental triplicates and are representative of at least three independent experiments. **P < 0.01; ***P < 0.001. BMDMs, bone marrow-derived macrophages; cCasp, cleaved caspase; cGSDM, cleaved gasdermin; h, hour; IL-1β, interleukin-1β; LDH, lactate dehydrogenase; LPS, lipopolysaccharide; min, minute; nig, nigericin; M-CSF, macrophage colony-stimulating factor; WT, wild type.

### CuET does not inhibit inflammasome-independent, GSDME-dependent pyroptosis

Raptinal and the combination of TNF-α and a TGF-β-activated kinase-1 (TAK-1) inhibitor both activate caspase-3 and GSDME-dependent pyroptosis independently of any inflammasome ^23, 68^. To determine whether CuET inhibits inflammasome-independent GSDME-maturation, BMDMs were treated with raptinal or TNF-α and TAK-1 inhibitor in the presence or absence of CuET. CuET did not inhibit the cleavage of GSDMD, GSDME, and caspase-3 induced by raptinal or TNF-α and TAK-1 inhibitor (Fig. 5A and B; Fig. S4A) and if anything appeared to enhance their cleavage in response to these stimuli (Fig. 5B). To determine why CuET inhibited the cleavage of these proteins in response to NLRP3 inflammasome activation but not apoptotic stimuli, we analyzed its effects on NF-κB and MAPK activation. CuET had no effect on LPS stimulated phosphorylation of IκBα, NF-κB/p65, and p38 MAPK (Fig. S4B). We also analyzed CuET effects on LPS plus nigericin inflammasome activation by measuring the formation of ASC specks, a key step in inflammasome assembly and activation, using WT BMDMs expressing fluorescent *Asc-citrine*. To minimize potential effects of CuET on inflammasome priming signals, the drug was added to LPS-primed cells and immediately followed by adding nigericin. The formation of ASC specks was significantly inhibited by CuET (Fig. 5C). These results suggest that CuET inhibits NLRP3 inflammasome activation, but not GSDME, caspase-3 or LPS-induced NF-κB or MAPK activation.

**Fig 5.**
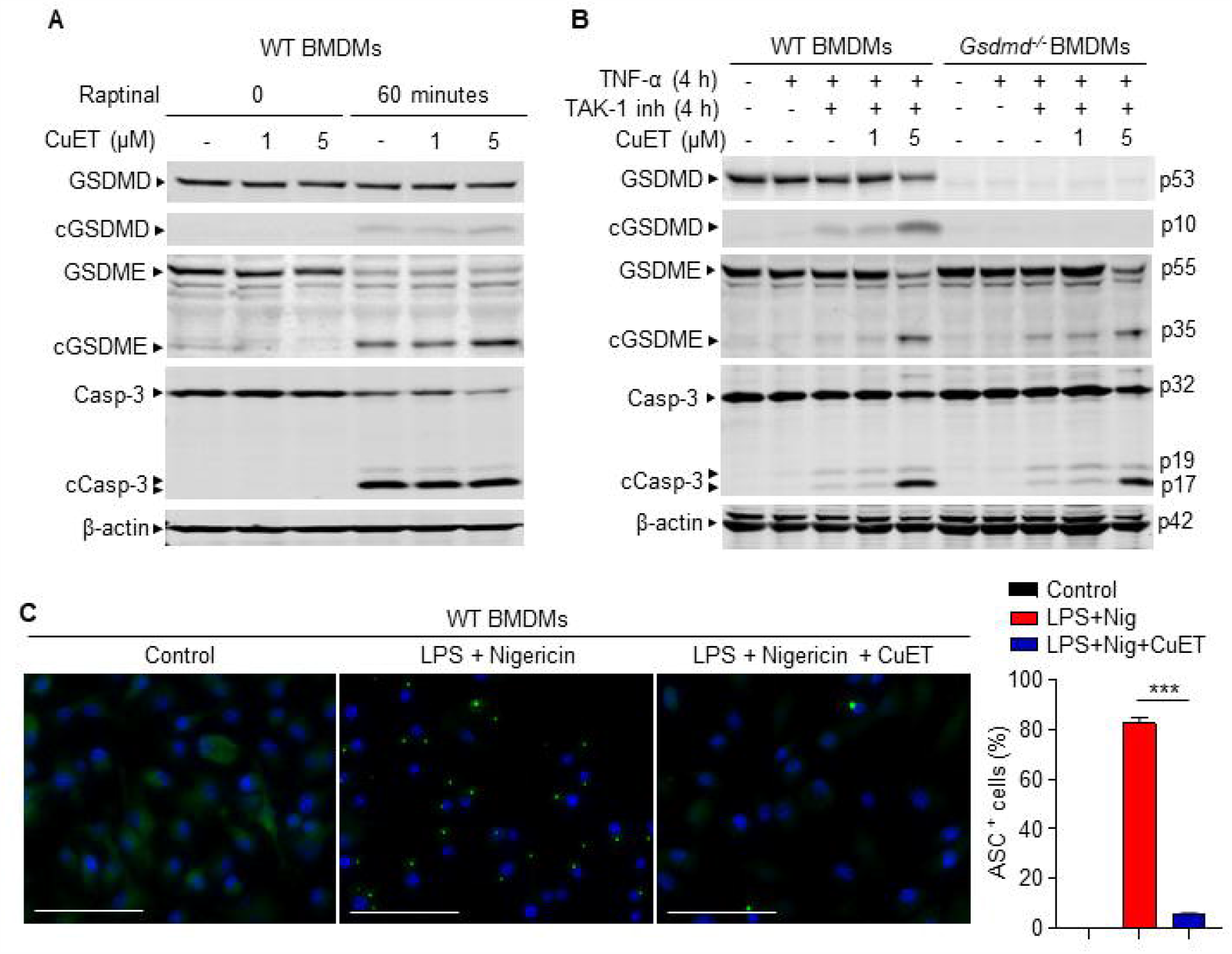
CuET inhibits NLRP3 inflammasome-driven ASC speck formation, but not inflammasome-independent processing of GSDMs and caspases. BMDMs were expanded *in vitro* M-CSF-containing media from bone marrow cells isolated from WT or *Gsdmd*^*−/−*^ mice. (A) WT BMDMs were pretreated with vehicle or CuET for 1 hour, and were stimulated with vehicle or 10 µM Raptinal for 1 hour. (B) WT and *Gsdmd*^*−/−*^ BMDMs were pretreated with vehicle or CuET for 1 hour, and were stimulated with 100 ng/ml TNF-α and 1 µM 5Z-7-oxozeaenol (TAK1 inhibitor) for 4 hours. Proteins in the whole cell lysates were analyzed by immunoblotting. (C) WT BMDMs from *Asc-citrine* mice were primed with 100 ng/ml LPS for 3 hours and treated with vehicle or 1 µM CuET for 15 minutes, then with15 µM nigericin for additional 30 minutes. ASC specks were visualized under fluorescence microscopy and quantified using imageJ. Scale bar, 25 μm. Data are mean ± SEM from experimental triplicates and are representative of at least three independent experiments. ***p < 0.001. BMDMs, bone marrow-derived macrophages; cCasp, cleaved caspase; CuET, bis(diethyldithiocarbamate)-copper; cGSDM, cleaved gasdermin; h, hour; IL-1β, interleukin-1β; LDH, lactate dehydrogenase; LPS, lipopolysaccharide; nig, nigericin; M-CSF, macrophage colony-stimulating factor; WT, wild type.

### CuET prevents the pathogenesis of mouse NLRP3^ca^ inflammasomopathy

Based on its in vitro potency in inhibiting the release of cytokines and pyroptosis, CuET should, in theory improve the disease outcomes of *Nlrp3*^*ca*^ mice. To test this idea, CuET was intraperitoneally injected to 10-day-old pups, once every 2 days ^69^. CuET significantly improved the survival rate of *Nlrp3*^*ca*^ mice (Fig. S5A) and reduced splenomegaly in these mice (Fig. S5B). Despite improved survival, treated-*Nlrp3*^*ca*^ mice still showed early mortality, likely due to the severe inflammation already activated before treatment was initiated. To control the onset of inflammation, we leveraged the *Nrlp3*^*+/fl(D301N)*^;*Cre-ER*^*TM*^ model for postnatal inducible activation of the NLRP3^ca^ (iNLRP3^ca^) inflammasome to test the efficacy of prophylactically administrated CuET. Three-month-old mice were injected with tamoxifen 3 times/week for 2 weeks. Vehicle or CuET were injected 2 days before starting tamoxifen and continued 3 times/week for 6 weeks. *iNlrp3*^*ca*^ mice all survived but developed cachexia, splenomegaly, leukocytosis and neutrophilia, which were significantly attenuated by CuET (Fig. 6A). CuET prevented inflammation in the liver and the disorganization of splenic architecture in *iNlrp3*^*ca*^ mice as assessed by H & E staining and had no apparent effect on these tissues in WT controls (Fig. 6B; Fig. S5C). Thus, the severe systemic multi-organ damage of *iNlrp3*^*ca*^ mice is attenuated by CuET.

**Fig 6.**
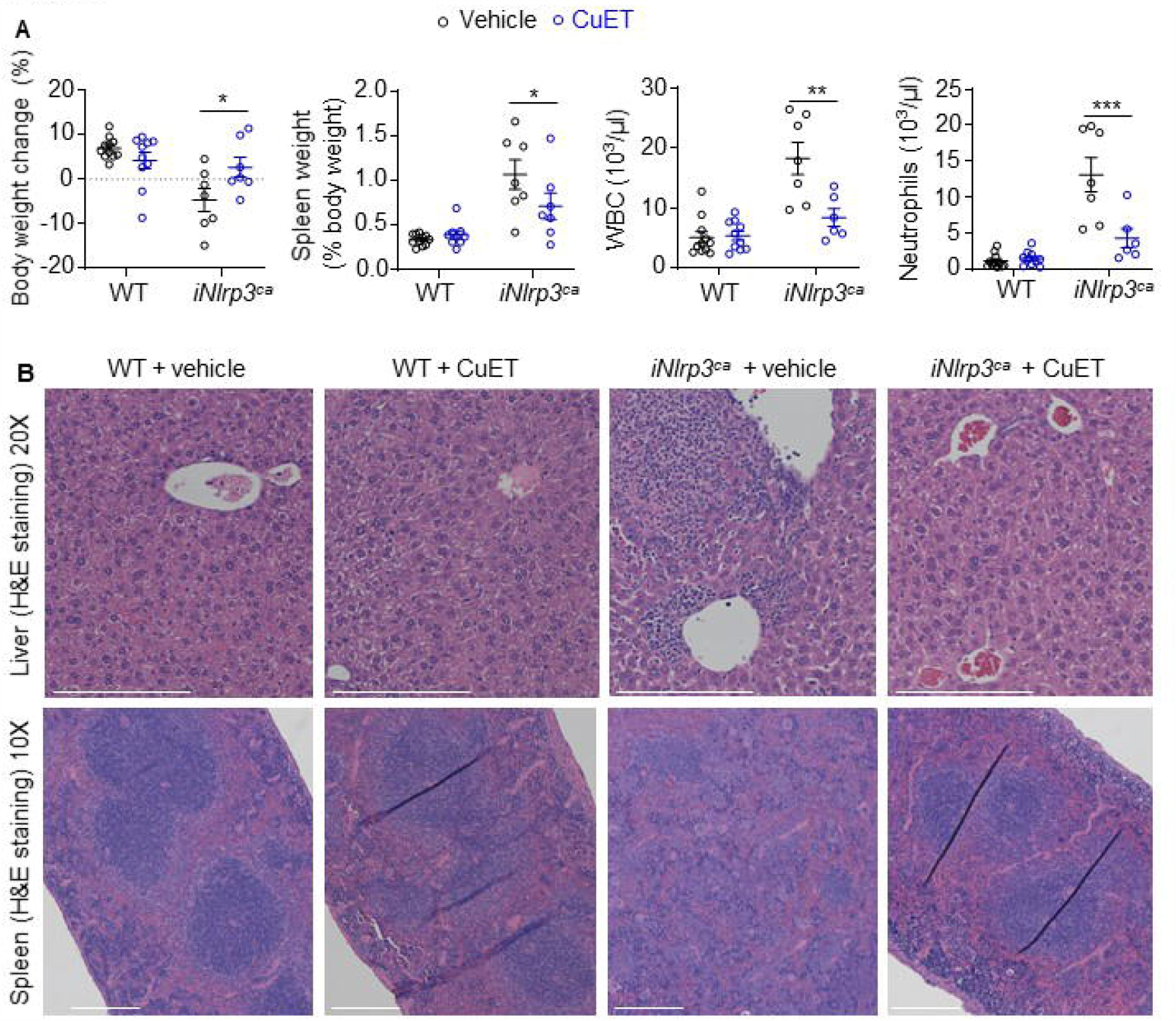
CuET prevents the pathogenesis of inflammasomopathy in the inducible NLRP3^ca^ (iNLRP3^ca^) model. Three-month-old mice were injected with tamoxifen once every other day, 3 times a week for 2 weeks. Injections with vehicle or 1 mg/kg body weight CuET started 2 days before the tamoxifen regimen, and were carried out once every other day, 3 times a week for 6 weeks. All injections were given intraperitoneally. (A) Body weight change, spleen weight, WBC; and neutrophils. Data are means ± SEM. *P < 0.05; **P < 0.01; ***P < 0.001. (B) H&E staining of liver and spleen sections. Scale bars, 200 μm. WBC, white blood cell; WT, wild-type; ca, constitutive activation; CuET, bis(diethyldithiocarbamate)-copper; iNLRP3^ca^, inducible NLRP3^ca^.

## Discussion

In contrast to *Nlrp3*^*ca*^ (NOMID) mice who die perinatally because of severe systemic multi-organ complications, *Nlrp3*^*ca*^ mice lacking GSDMD (*Nlrp3*^*ca*^;*Gsdmd*^*-/-*^ mice) grow and age normally and are fertile. Consistent with these observations, ablation of GSDMD reduces inflammation and improves disease outcomes in mouse models of sepsis, Familial Mediterranean Fever (FMF), and experimental autoimmune encephalitis ^14, 55, 70^. However, when stressed by exposure to exogenous inflammatory stimuli (sublethal doses of LPS or TNF-α), *Nlrp3*^*ca*^;*Gsdmd*^*-/-*^ mice unexpectedly secreted high amounts of IL-1β and IL-18. The combination of these inflammatory stimuli with constitutively active NLRP3 overwhelmed the protection provided by GSDMD or caspase-1 deficiency, rewiring signaling cascades through caspase-3 and GSDME to cause pyroptosis ^22, 25^. These observations suggest that health complications such as those associated with infection or inflammatory comorbidities have the potential to exacerbate inflammation driven by dysregulated inflammasomes, potentially limiting the efficacy of GSDMD-based therapies. This scenario can be tested by assessing the consequences of infections or injuries in NOMID or FMF GSDMD deficient mice, which can be viewed as animals with an underlying inflammatory condition.

Both *Nlrp3*^*ca*^;*Gsdmd*^*-/-*^ BMDMs as well as *Gsdmd*^*-/-*^ BMDMs produced IL-1β and underwent pyroptosis in response to LPS, although these responses were delayed in both cell lines compared to WT cells, occurring only upon sustained exposure to NLRP3 inflammasome activators. The responsiveness of *Gsdmd*^*-/-*^ cells to LPS *in vitro* suggests that exposure of GSDMD-deficient mice to endotoxin for a period longer than the 6-hour time-point of this study may lead to significant IL-1β and IL-18 secretion. Consistent with this, a small proportion of *Gsdmd*^*-/-*^ mice die upon LPS-challenge ^14, 62^. Pyroptosis and cytokine release by GSDMD or caspase-1 deficient cells strongly correlated with caspase-3 and GSDME activation and were significantly suppressed in *Gsdmd*^*-/-*^;*Gsdme*^*-/-*^ cells, suggesting that caspase-3 activation of GSDME was largely responsible for inflammatory death when NLRP3 is activated but the canonical caspase-1-GSDMD pathway is not available. Consistent with our findings, a recent study reported that the TLR1/TLR2 agonist Pam3CSK4 promotes the release of IL-1α by caspase-1 deficient macrophages through GSDME conduits ^28^. In our study, caspase-3 activation was enhanced by GSDMD or caspase-1 genetic deficiency and by GSDMD inhibition with CuET. We did not explore the mechanism responsible for increased caspase-8 and caspase-3 activation in these settings, but one possibility is that caspase-8 is recruited to the NLRP3 inflammasome to some extent even when caspase-1 is present - or more so when it is not - and that inflammasome-bound caspase-8 is very efficient at processing caspase-3, which in turn activates GSDME. It is worth noting that there was some residual IL-1β release and pyroptosis even in *Gsdmd*^*-/-*^;*Gsdme*^*-/-*^ BMDMs, suggesting that another GSDMD and GSDME independent inflammation pathway can be activated downstream of NLRP3, potentially involving another caspase and/or another gasdermin. From our study it remains uncertain whether cytokines are secreted through GSDME pores in the setting of caspase-1 or GSDMD deficiency or are passively released when the cell membrane is grossly disrupted and LDH is released, as was recently proposed for *in vitro* GSDMD-mediated HMGB1 release ^71^. Taken together, these results suggest that macrophages have more than one salvage mechanism to ensure that signals transmitted by inflammasome activators result in inflammation.

Drug discovery and repurposing efforts have identified the FDA-approved drugs dimethyl fumarate and disulfiram as GSDMD inhibitors. Covalent modification of Cys191 in human and Cys192 in mouse is the proposed main mechanism through which disulfiram inhibits GSDMD pore-forming activity, although dimethyl fumarate likely works through another mechanism ^62^. Here, we report that the disulfiram metabolite CuET exhibits remarkable efficacy in the *Nlrp3*^*ca*^ inflammasomopathy model. Mechanistically, we find that CuET suppresses cytokine secretion and pyroptosis caused by both GSDMD and GSDME. However, CuET inhibits GSDME cleavage and pyroptosis in cells stimulated with NLRP3 inflammasome activators but not apoptotic stimuli. These results rule out GSDME as the major direct target of CuET inhibitory actions, a view that is consistent with the compound’s ability to inhibit the formation of ASC specks. Although, as far as we are aware, direct inhibition of GSDME pore formation by disulfiram has never been measured, Cys191/192 is not conserved in mouse GSDME and therefore disulfiram is not predicted to be a potent regulator of GSDME. However, as a cysteine-reactive drug, disulfiram has the potential to inactivate other enzymes, including the caspase cysteine proteases and other proteins with reactive cysteines that are modulated by cellular redox status, which include the NLRP3 inflammasome ^72^. When a potent drug target such as GSDMD is not present, disulfiram may switch to inhibit other less reactive substrates. Further work is needed to investigate how CuET inhibits ASC polymerization and whether it inhibits other early steps of inflammasome signaling.

This study reveals that ablation of GSDMD in *Nlrp3*^*ca*^ mice does not prevent these animals from producing excessive levels of IL-1β and IL-18 in response to inflammatory challenges (Fig. 7). Importantly, the disease severity in *Nlrp3*^*ca*^ mice was remarkably reduced by CuET, a drug that interferes directly or indirectly with both GSDMD-dependent and independent inflammation. Our findings suggest that disulfiram might be worth testing in CAPS patients for whom existing therapies that inhibit IL-1 or other inflammatory cytokines do not adequately suppress disease symptoms.

**Fig 7.**
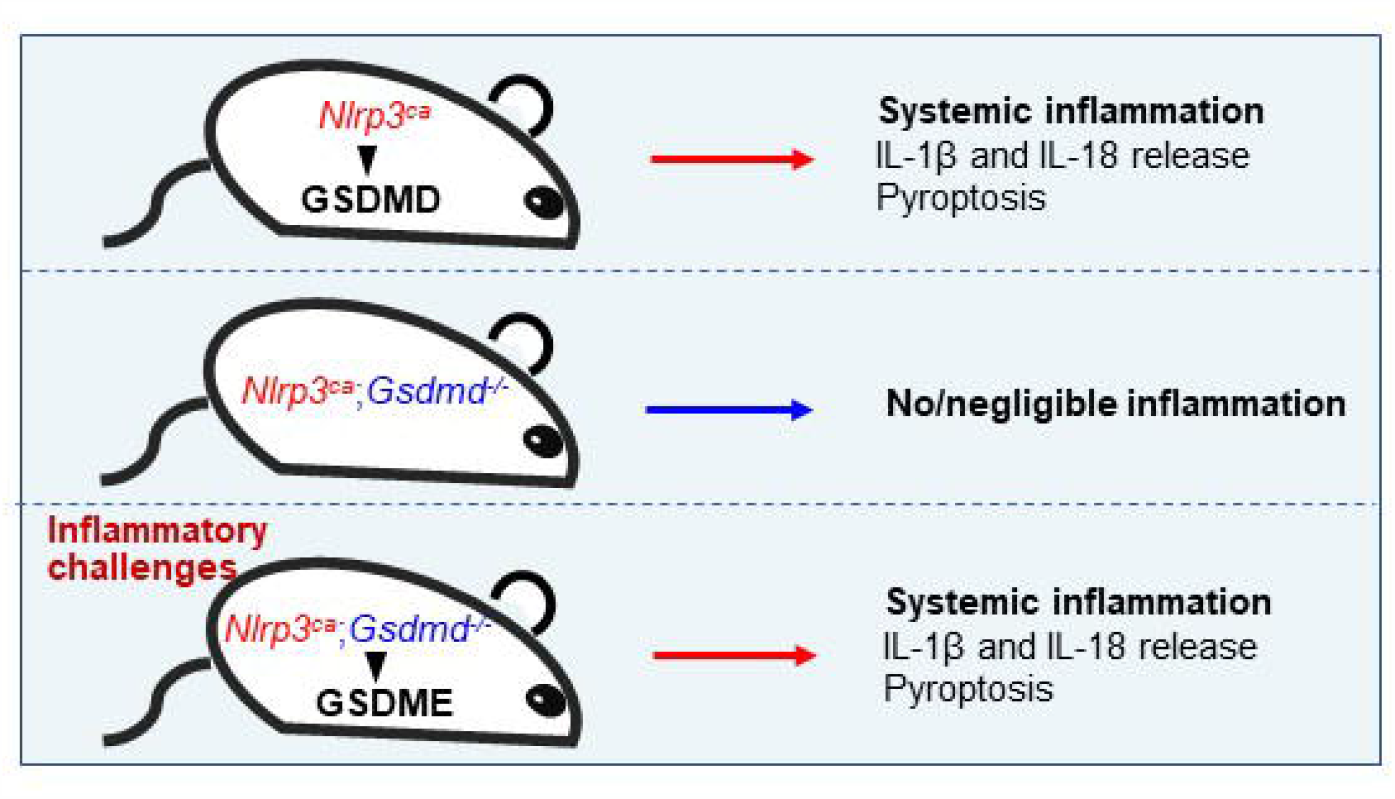
A model showing that inflammatory challenges activate GSDME to reverse the beneficial actions of GSDMD deficiency in cells expressing NLRP3^ca^. The NLRP3^ca^ inflammasome promotes inflammation through GSDMD as inflammation is negligible in *Nlrp3*^*ca*^;*Gsdmd*^*−/−*^ mice. However, these compound mutant mice develop systemic inflammation through GSDME when exposed to inflammatory challenges.

## Materials and methods

### Animals

*Casp1*^*−/−*^, *Cre-ER*^*TM*^ (B6.Cg-Tg(CAG-cre/Esr1*)5Amc/J) mice, *lysozyme M-Cre* mice and *Nlrp3fl*^*(D301N)/+*^ mice have been previously described ^50, 52, 73^. *Nlrp3*^*ca*^ mice with constitutive activation of NLRP3 in myeloid cells driven by *lysozyme M-Cre* have been previously described ^52^. *Cre-ER*^*TM*^ mice and *Nlrp3*^*fl(D301N)/+*^ mice were crossed to generate *Nlrp3*^*fl(D301N)/+*^;*Cre-ER*^*TM*^ mice and *Cre-ER*^*TM*^ mice. Injection of tamoxifen (i.p., 75 mg/kg body weight; Sigma-Aldrich) to *Nlrp3*^*fl(D301N)/+*^;*Cre-ER*^*TM*^ mice and *Cre-ER*^*TM*^ mice to yield inducible *Nlrp3*^*ca*^ (*iNlrp3*^*ca*^) mice and control mice, respectively, has been previously described ^73^. *Gsdmd*^−/−^ mice were kindly provided by Dr. V. M. Dixit (Genentech, South San Francisco, CA). *Asc-citrine* and *Gsdme*^*-/-*^ mice were purchased from The Jackson Laboratory (Sacramento, CA). All mice were on the C57BL6J background. All procedures were approved by the Institutional Animal Care and Use Committee of Washington University School of Medicine in St. Louis.

### Administration of drugs, LPS, and TNFα

Mice were i.p. injected with CuET (TCI America, OR) (1 mg/kg) formulated in sesame oil (0.2 mg/ml) or vehicle, once every other day, 3 times per week. Ten day-old *Nlrp3*^*ca*^ mice were treated with vehicle or CuET for 9 weeks; 12-week-old *iNlrp3*^*ca*^ and control mice were treated with vehicle or CuET 3 times/week for 6 weeks. For LPS challenge experiments, WT, *Gsdmd*^*-/-*^, *Gsdme*^*-/-*^, *Gsdmd*^*-/-*^;*Gsdme*^*-/-*^, and *Nlrp3*^*ca*^;*Gsdmd*^*-/-*^ mice were i.p. injected with 15 mg/kg LPS (*Escherichia coli* O111:B4, Sigma, MO). Mice were i.p. injected with 0.5 mg/kg TNFα (Biolegend, CA). Blood was collected by cardiac puncture 6 hours or 2 hours after LPS or TNFα challenge, and was allowed to clot at room temperature. Serum obtained after centrifugation at 2,000 x g for 10 minutes was used for determinations of cytokine and chemokine levels by V-PLEX Plus Proinflammatory Panel1 Mouse Kit (Meso Scale Diagnostics, MD), except IL-18, which was analyzed by enzyme linked immunosorbent assay (ELISA) kit (Sigma, MO).

### Cell cultures

BMDMs were obtained by culturing mouse bone marrow cells in culture media containing a 1:10 dilution of supernatant from the fibroblastic cell line CMG 14-12 as a source of macrophage colony-stimulating factor ^74^, a mitogenic factor for BMDMs, for 4-5 days in a 15-cm dish as previously described ^73^. Briefly, nonadherent cells were removed by vigorous washes with PBS, and adherent BMDMs were detached with trypsin-EDTA and cultured in culture media containing a 1:10 dilution of CMG for various experiments.

For all *in vitro* pharmacology experiments except otherwise specified, cells were pre-treated with vehicle (0.1% DMSO, final concentration) or inhibitors (in 0.1% DMSO, final concentration) for 1 hour before stimulation with the indicated ligand or ligands. Protein expression was analyzed by ELISA or Western blot. To activate the NLRP3 inflammasome, BMDMs were plated at 10^4^ cells per well on a 96-well plate or 10^6^ cells per well on a 6-well plate overnight. Cells were primed with LPS and then with 15 μM nigericin (AdipoGen) as indicated, and conditioned media were collected for the analysis of IL-1β and LDH.

### Histology

Tissue samples were processed as described previously ^73^. In brief, liver, and spleen were fixed in 10% formalin overnight. Fixed tissues were embedded in paraffin, sectioned at 5-μm thicknesses, and mounted on glass slides. The sections were stained with H&E as described previously ^73^. Photographs were taken using ZEISS microscopy (Carl Zeiss Industrial Metrology, MN).

### Peripheral blood analyses

Complete blood counts were performed by the Washington University School of Medicine as previously described ^52^.

### Western blot analysis

Cell extracts were prepared by lysing cells with RIPA buffer (50 mM Tris, 150 mM NaCl, 1 mM EDTA, 0.5% NaDOAc, 0.1% SDS, and 1.0% NP-40) plus phosphatase and protease inhibitors (2 mM NaVO4, 10 mM NaF, and 1 mM PMSF) and Complete protease inhibitor cocktail (Roche, CA). For tissue extracts, liver tissues were homogenized and lysed with RIPA buffer containing phosphatase and protease inhibitors. Protein concentrations were determined by the Bio-Rad Laboratories method, and equal amounts of proteins were subjected to SDS-PAGE gels (12%) as previously described ^54^. Proteins were transferred onto nitrocellulose membranes and incubated with antibodies against GSDMD (1;1,000; Abcam, MA), GSDME (1;1,000; Abcam, MA), caspase-1 (1;1,000; Abcam, MA), caspase-3 (1:1,000; Cell Signaling Technologies, MA), p38 MAPK (1:1,000; Cell Signaling Technologies, MA), β-actin (1:2,000; Santa Cruz Biotechnology, TX), p65/p-p65 (1:1,000; Cell Signaling Technologies, MA), IκBα/p-IκBα (1:1,000; Cell Signaling Technologies, MA) overnight at 4°C followed by incubation for 1 hour with secondary goat anti–mouse IRDye 800 (Thermo Fisher Scientific, MA) or goat anti–rabbit Alexa Fluor 680 (Thermo Fisher Scientific, MA), respectively. The results were visualized using the Odyssey infrared imaging system (LI-COR Biosciences, NE).

### LDH assay and IL-1β ELISA

Cell death was assessed by the release of LDH in conditioned medium using LDH Cytotoxicity Detection Kit (TaKaRa, CA). IL-1β levels in conditioned media were measured by ELISA (eBiosciences, NY).

### ASC specks assay

*Asc-citrine* BMDMs were plated at 10^4^ cells per well on a 16-well glass plate overnight. Cells were primed with LPS for 3 hours and pretreated or not with CuET for 15 minutes before adding 15 μM nigericin (AdipoGen, CA) for 30 minutes. Cells were washed with PBS and fixed with 4% polyformalin buffer for 10 minutes at room temperature. Photographs were taken using ZEISS microscopy (Carl Zeiss Industrial Metrology, MN). Quantification of ASC specks was carried out using ImageJ.

### Statistical analysis

Statistical analysis was performed using the Student’s t test, one-way ANOVA with Tukey’s multiple comparisons test, or two-way ANOVA with Tukey’s multiple comparisons test as well as the Log-rank (Mantel-Cox) test for comparison of survival curves in GraphPad Prism 8.0 Software.

## Acknowledgements

This work was supported by NIH/NIAMS AR068972 and AR076758 grants to G.M. Y.A-A. was supported by NIH grants AR049192, AR054326, AR072623 and by a grant #85160 from the Shriners Hospital for Children.

We want to thank Dr. Deborah J. Veis for critical reading of this manuscript.

