## Supplemental Figure legends for "Activation of GSDME compensates for GSDMD deficiency in a mouse model of NLRP3 inflammasomopathy"

**Supplemental Figures**

**Fig. S1**. GSDMD deficient mice expressing hyper-activated NLRP3 inflammasome age normally. (A) Survival, WBC, and percent of spleen weight of *Gsdmd^−/−^*, *Nlrp3^ca^*, and *Nlrp3^ca^*;*Gsdmd^−/−^* mice. Data are mean ± SEM. Dashed lines delineate normal ranges. (B and C) Representative images of liver and spleen from 6- and 12-month old mice. (C) High magnification of the boxed area. Scale bars, 200 µm. WBC, white blood cell; GSDMD, gasdermin D; ca, constitutive activation.

**Fig. S2**. LPS and nigericin induce the release of IL-1β and LDH by *Gsdmd^−/−^* but not *caspase-1^-/-^* BMDMs. (A, B, and D) *In vitro*-expanded BMDMs from WT and *Gsdmd^−/−^* mice were primed with LPS for 3 hours and treated with nigericin for 2 hours at the indicated concentrations. (C and E) BMDMs from *caspase-1^−/−^* mice were primed with LPS for 3 hours and treated with 15 µM nigericin for 20 minutes, 40 minutes, 1, 2, 3, 4 hours. Proteins in the whole cell lysates were analyzed by immunoblotting. IL-1β and LDH in the conditioned media were measured by ELISA and by the cytotoxicity detection Kit, respectively. Dash lined boxes were drawn to fill the void. Data are mean ± SEM from experimental triplicates and are representative of at least three independent experiments. BMDMs, bone marrow-derived macrophages; ca, constitutive activation; cCasp, cleaved caspase; h, hour; IL-1β, interleukin-1β; LDH, lactate dehydrogenase; LPS, lipopolysaccharide; M-CSF, macrophage colony-stimulating factor; WT, wild type.

**Fig. S3**. CuET inhibits the remaining of IL-1β secretion and pyroptosis in *caspase-1^-/-^* BMDMs and is as potent as disulfiram. (A and B) *In vitro*-expanded BMDMs from *caspase-1^-/-^* mice were primed with LPS for 3 hours and treated with vehicle or CuET for 1 hour before adding 15 µM nigericin for 3 hour. (A) Proteins in the whole cell lysates were analyzed by immunoblotting. (B) IL-1β and LDH in the conditioned media were measured by ELISA and by the cytotoxicity detection Kit, respectively. WT BMDMs were primed with LPS for 3 hours and treated with CuET (C and D) or disulfiram (E and F) for 1 hour at the indicated concentrations, then with nigericin for additional 1 hour. IL-1β and LDH in the conditioned media were measured by ELISA and by the cytotoxicity detection Kit, respectively. Data are mean ± SEM from experimental triplicates and are representative of at least three independent experiments. ***P < 0.001. BMDMs, bone marrow-derived macrophages; cGSDMD, cleaved GSDMD; CuET, bis(diethyldithiocarbamate)-copper; DSF, disulfiram; h, hour; IL-1β, interleukin-1β; LDH, lactate dehydrogenase; LPS, lipopolysaccharide; Nig, nigericin; WT, wild type.

**Fig. S4**. CuET does not inhibit inflammasome priming signals and apoptotic stimuli-induced maturation of GSDMs and caspases. (A) WT BMDMs were pretreated with vehicle or CuET for 1 h, and were stimulated with 10 µM Raptinal for 1 or 1.5 hour. (B) WT BMDMs were pretreated with vehicle or CuET for 1 hour and stimulated with LPS for 30 minutes or 1 hour. Proteins in the whole cell lysates were analyzed by immunoblotting. BMDMs, bone marrow-derived macrophages; ca, constitutive activation; cCasp, cleaved caspase; cGSDME, cleaved DSME; IL-1β, interleukin-1β; LPS, lipopolysaccharide; min, minute; nig, nigericin; p38, p38 MAPK; WT, wild type.

**Fig. S5**. CuET improves the pathogenesis of *Nlrp3^ca^* mice. Ten-day old *Nlrp3^ca^* mice were injected intraperitoneally with vehicle or 1 mg/kg body weight CuET for 9 weeks. (A) Survival (12-13 mice/group) was scored. (B) Spleen weights of survived mice. Data are mean ± SEM from 2-4 mice/group. *P < 0.05; ***P < 0.001. (C) Three-month-old WT and *iNlrp3^ca^* littermate mice were treated with CuET (1 mg/kg body weight) or vehicle by intraperitoneal injection for 6 weeks, starting 1 day before tamoxifen administration for 2 weeks. H&E staining of liver and spleen sections. Scale bars, 200 μm. ca, constitutive activation; CuET, bis(diethyldithiocarbamate)-copper; iNLRP3^ca^, inducible NLRP3^ca^; WT, wild-type.
