## Supplementary figures and images for "Activation of GSDME compensates for GSDMD deficiency in a mouse model of NLRP3 inflammasomopathy"

### Supplemental Figure 1

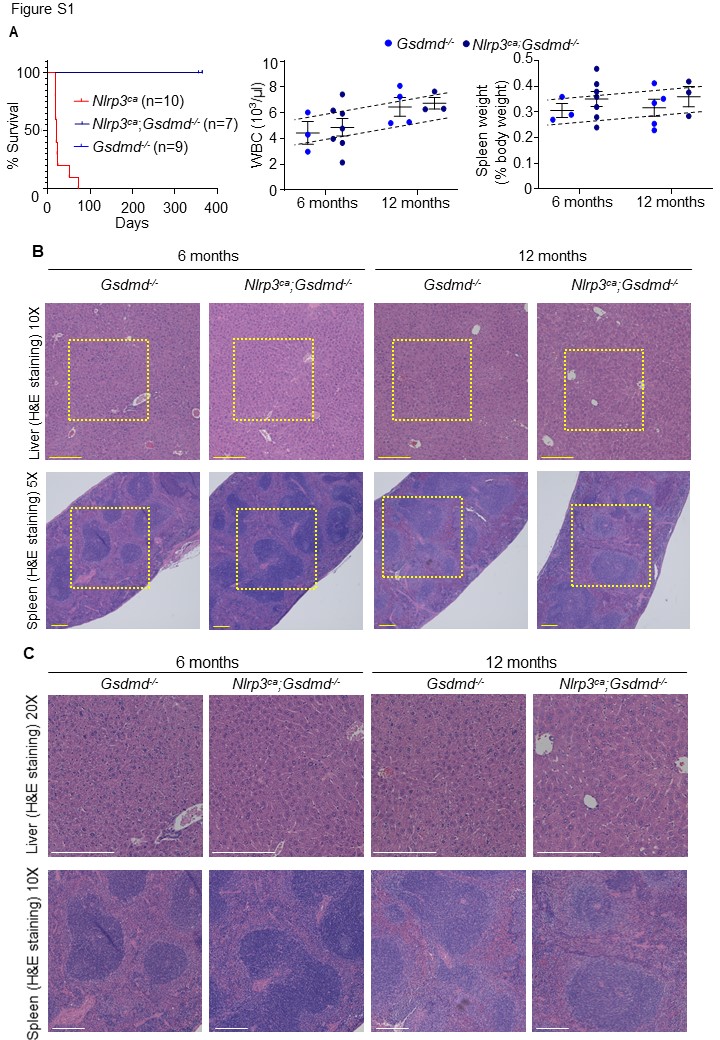

### Supplemental Figure 2

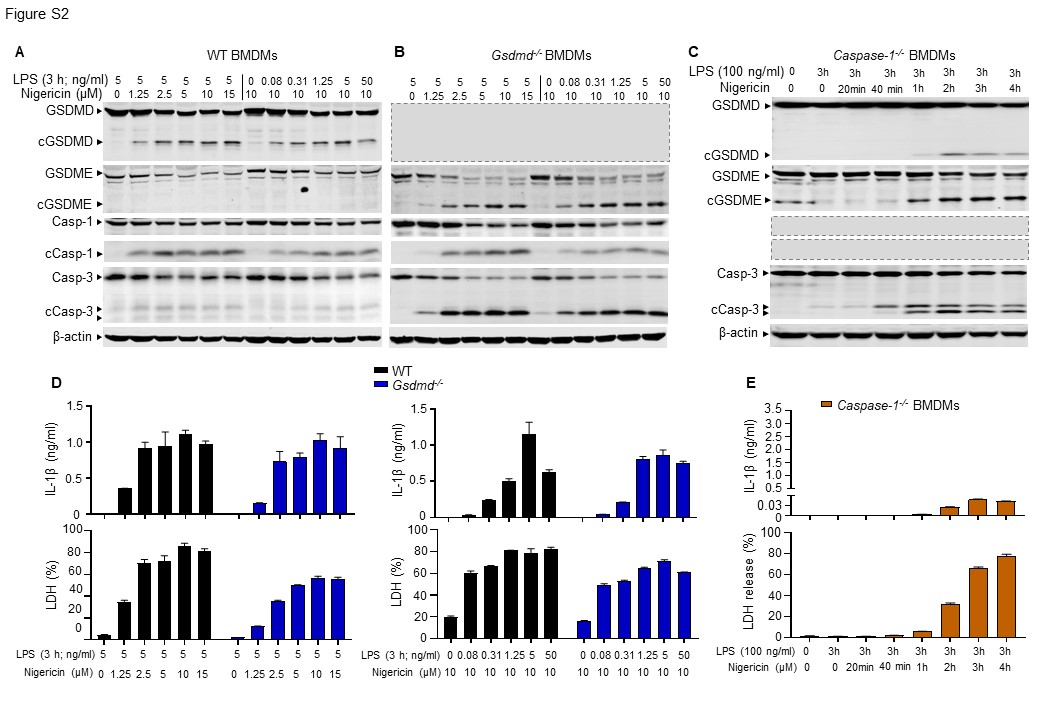

### Supplemental Figure 3

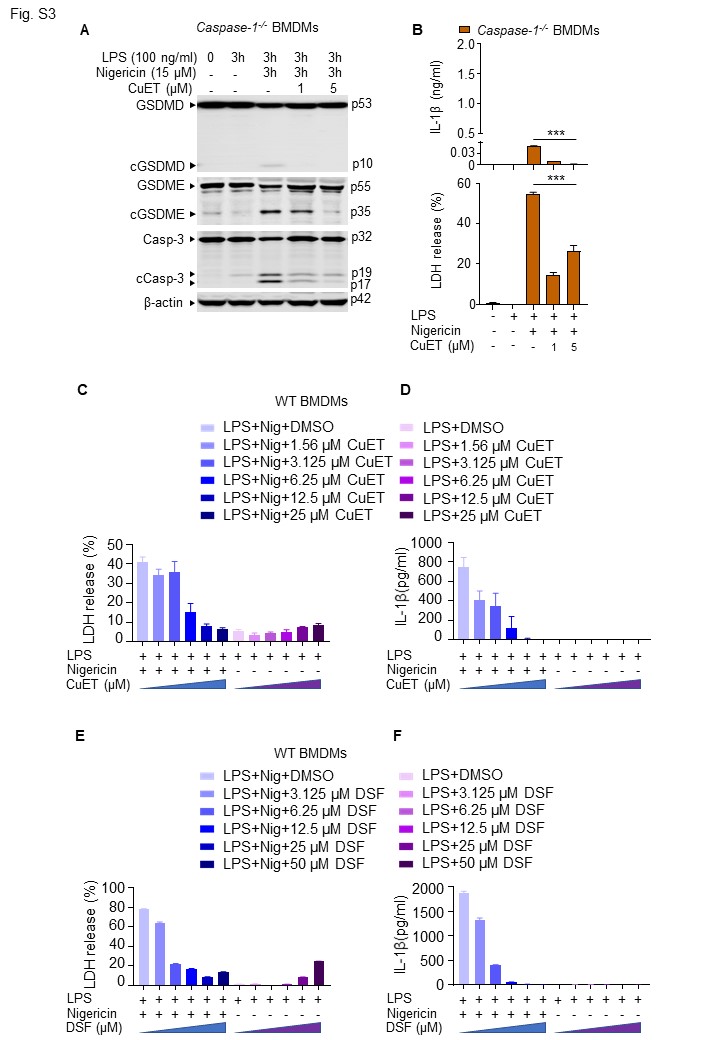

### Supplemental Figure 4

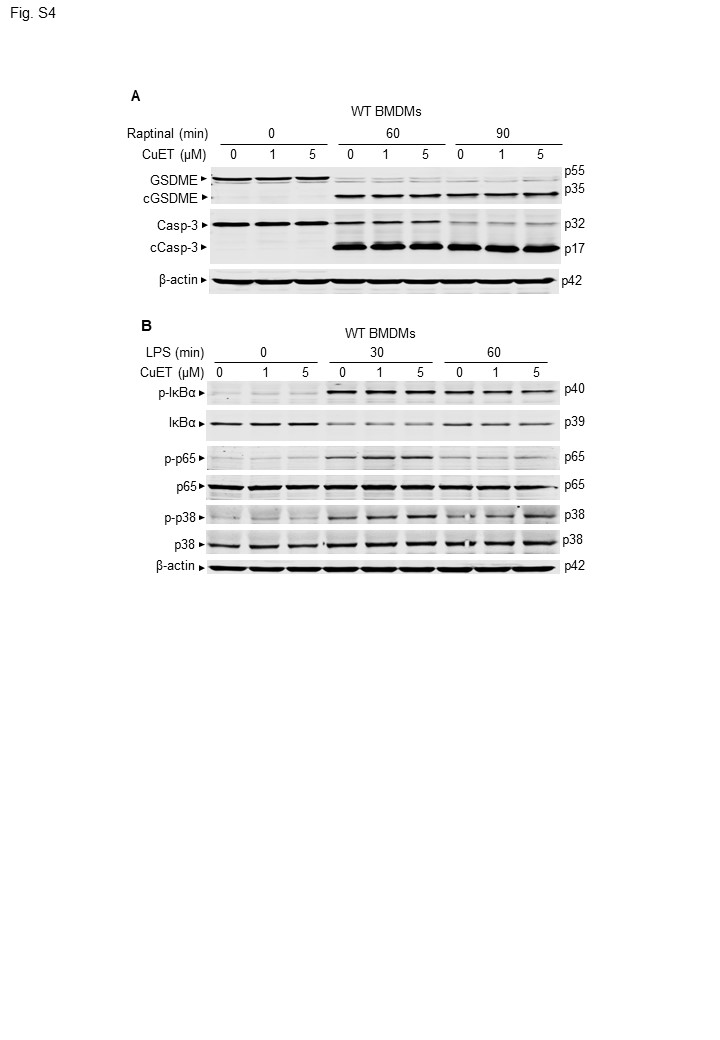

### Supplemental Figure 5

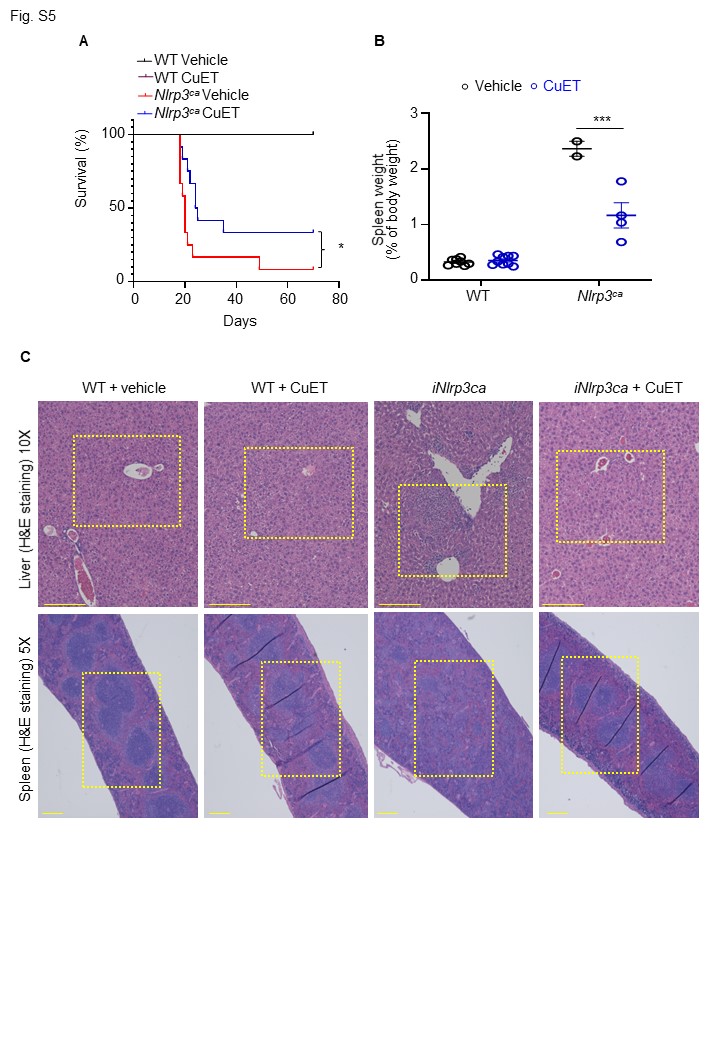
